# Modeling of Nano-carriers for Vascular-targeted Delivery for Blood Clots Treatment

**DOI:** 10.1101/2020.07.02.184242

**Authors:** Ibtihag Yahya, Megdi Eltayeb

## Abstract

Nanoparticles have become one of the emerging and promising technologies that revolutionized the medical field’s future on which has received much attention from the scientific community and researchers. Nanotechnology-based targeted drug delivery has a high capacity for loading large amounts of anticoagulants drug to dissolve clots in a safe manner without affecting healthy blood vessels. This paper seeks to provide a better understanding of both the anticoagulant drug release process and the coagulation eliminating process by simulating each process using chemical reaction engineering, moving mesh, and convection-diffusion equation modules. This study adds to a growing corpus of research showing that nanotechnology empowers in treating blood clots within 2-4 hours. In addition, these results cast a new light on a better understanding of the anticoagulation drug diffusion from both spheres and multiply-twinned nanoparticles besides the reduction of clots growth and how it dissolved over time.

## 1. Introduction

The nanoparticle-based drug delivery researches have a long tradition. Today, nanotechnology become at the forefront of the medical research basics that are attracting many researcher’s attention. There are growing appeals for investigating the use of nanotechnology in the treatment and prevention of the blood clot. As a rule, thrombosis is a term used to express the blood clots or blood clump that is caused due to the blood transformation from a liquid state to a semi-solid or gel-like state [1]. The sudden interruption of blood flow due to travel of blood clot (embolus) further down the arteries that become stuck and adhered to the artery wall and then; prevents the flow of blood to an organ or part of the body known as an “arterial embolism”.

To demonstrate, both nanoparticles and polymers can be integrated into many materials and applications such as biotechnology, pharmaceuticals, protective coatings, chemical catalysis and the most important of which is drug delivery [2,3]. Therefore, one of the most important basics that must be known and considered when using nanoparticles in particular for medical applications is the ability to determine their chemical and physical properties, and this can only be done through an understanding and knowledge of the morphological stability of these particles [4]. Moreover, the physical and chemical properties of nanomaterials such as size, composition, shape, and charge determine the interaction of these materials with the different components of the coagulation system [5]. Whereas, manipulating the properties of nanoparticles during the formulating process is an important rule that assists in providing a deep understanding of the drug release profiles from the beginning of the process until this medication delivered to the desired location [6–8]. E.g., the nanoparticles can be engineered so that it can interact with the components of the clotting system to remove it within hours to avoid complications such as autoimmune reactions and deaths due to transmission of this thrombus to the lung or heart in addition to the occurrence of cancers and cardiovascular disorders [9,10].

Indeed, one of the problems with traditional methods of treating blood clots is the drug ineffectiveness in removing the blood clot within hours, which leads to preventing blood flow inside the blood vessels and then the death of tissue because the insufficient blood and oxygen that reach them. In addition to the large dose of medication provided to the patient.

Most of the theories of blood clot removal are however focused on explaining the methods to treat, prevent blood clots, their signs, symptoms, causes as well as the available drugs of treatments and its side effects. In the past few decades, nanotechnology for drug delivery systems and drug-release kinetics have played an important role in the medical field especially in the diagnosis and treatment of various diseases such as cancer detection, targeting, DNA delivery and the biochemical sensors [11,12]. A study was held at Harvard University’s Wyss Institute to treat the coagulation disorders using nanoparticles, where it secretes the drug in the place of clotting when it is exposed to high shear pressure due to the narrowing of the blood vessels due to stroke [13]. Other researchers [14,15] have used magnetic nanoparticles to control the medication to dissolve the clot. The study concluded that this mechanism is highly effective and effective in dissolving clots within hours safely. A clinical study conducted to suggested a classification system to predicts the clot type, number of affected segments, thickness, area and circumference with a scale from 0-10 [16]. This study proved the validity of this system. Few studies have examined the multiple twinning nanoparticle shapes, despite its widespread nature, in various crystalline materials. Two papers were published providing an overview of the growth of nanoparticles, their applications, characterization, properties, structural and thermal stabilities, and focused on highlighting the comparison of both single-crystal particles versus double Multiply-twinned particles in order to study controlled growth and determine the differences between them [17,18].

Despite all researches that have been done; still, there are no studies have focused on the drug release kinetics simulation in the blood clot site, in addition, to examine the drug release rate as well as the thrombosis elimination over time and this is what this study attempting to shows. Also, there are no studies that focused specifically on investigating the diffusion of the drug from different forms of particles, and this is what the study will achieve for both the spherical morphology and the multiple-twinned of the prism-shape to determine which is optimal for application in the treatment of blood clots. So, this paper considers and focuses on the arterial blood clot (thrombosis) and we simulated the antithrombotic agent delivery and release to the blocked artery in the artery using chemical reaction engineering, moving mesh and convection-diffusion equation modules as the main subject of its study to provide a clear view of the dissolution and treatment of blood clots.

## 2. Theory

In fact, the clotting process is a double-edged sword, while it is a necessary life-saving process, especially when it comes to preventing excessive bleeding when a blood vessel is injured [19], on the other hand, it can be life-threatening and fatal when this clot becomes immoderate blood clot (Hypercoagulation/Thrombotic disorders) [20] and causes many diseases such as heart attacks, strokes, kidney failure, pulmonary embolism, arterial thrombosis, deep vein thrombosis (DVT), venous thromboembolism (VTE), peripheral artery disease (PAD) and pregnancy-related problems [21–24].

Thrombosis in a vein or an artery is a blood clot formation which leads to limited or blocked blood flow due to narrowing of the artery channel and then the reduction of the oxygen and the nutrition and thus the death of the surrounding tissue [25]. To demonstrate, the thrombosis of an artery may cause a stroke in the brain or myocardial infarction in the heart [26]. The main three causes of thrombosis are vascular injury, stasis, and hypercoagulability [27]. The clot either be a thrombus that extends to the wall but does not close the blood vessel and is known as the mural thrombus. Either the thrombus blocks blood flow and this closes the blood vessel completely and is known as an occlusive thrombus [28]. As for the third type, it is a choice that extends and multiplies along the vessel and is known as reproductive thrombosis [29]. Arterial thrombosis can break off formulating a clot part called an embolism which is life-threating as it can be traveling to the lung or to the hart.

## 3. Materials and Methods

The polymers have effective applications in both delivery and release of drugs particularly in dissolving and removing clots [30,31]. Polymers have been used recently in many fields, especially medical due to the fact that it is one of the organic materials in addition to, its unique properties such as ease of manufacture, bio-compatible, and controllable structures during design [32,33]. In the drug delivery and given the small size of these polymeric nanoparticles basis their efficiency in controlling the drug and active agents release makes it ideal for all the medical applications. Moreover, its flexibility leads to enhancing the penetration of this drug into the target location and the ability to control its melting rates; it made it best possible for achieving the desired therapeutic goals with superior effectiveness [34,35].

This study focused on modeling the diffusion release behavior of a hybrid medicine consist of both ‘Tissue Plasminogen Activator’ (tPA) drug in a combination with an anti-clotting medicine (Heparin) to simulate and predict the arterial clots dissolving and removal through intra-arterial injection of polymer–colloid composite nanoparticles carriers in sizes ranging from 0.01-0.03 nm. In addition, evaluating the effective concentration of the drug needed inside the polymer–colloid particles. We classify the thrombosis using the classification system as showed in (Figure 1) this thrombosis has a smooth type with ≥ 6 mm thickness and with an area about 25-50% and 91-180° circumference [26]. The anticoagulant drug release after injected into the blood clot site and the coagulation eliminating by the interaction between polymeric nanoparticles and blood clots processes were simulated using different modules:

**Figure 1.**
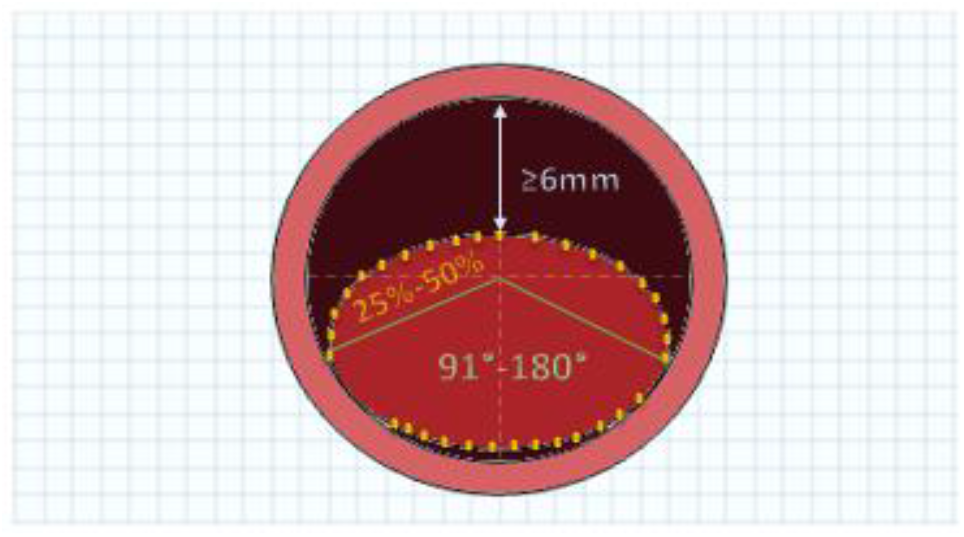
Cross-section of the Artery to classify the thrombus (type, thickness, area, and circumference).

### 3.1 Chemical Reaction Engineering Module

The modeling of chemical systems can be simulated in a close manner that mimics the reality by using the chemical reaction engineering module i.e., modeling the reaction kinetics, temperature, and flow of fluids, which are mainly affected by the chemical composition of materials and these systems are modeled as functions in space and time. As the nanoparticles are one of the most promising treatments for the blood clots. This model describes and simulates the release of hybrid anticoagulation that consist of both ‘Tissue Plasminogen Activator’ and ‘Heparin’ drugs to the site of the thrombus in a bloody artery, which are effective in disintegrate, remove blood clots and prevent the growth of a blood clot using 2D axial symmetrical geometry and a mirror plane.

This model consists of a blood clot, polymeric nanoparticles with sphere and prism-shapes, and an arterial vessel as shown in (Figure 2 A, B). The reaction kinetics of the drug is analyzed in two parts. The first part is represented in 0D in which the engineering interface for the reaction is used and allows the pharmacokinetics to be resolved to interact with the drug over time and describe the reaction system. As part of the drug is separated from the nanocarriers and the other remains attached to it. The nanoparticles decompose to release the drug due to the shear stress/ pressure and once the release is complete, the two types of medicine separate from each other. The second part, which is represented in 2D, which shows how the drug moves in the place of the blood clot in order to track the movement of the drug particles. Where it is examined how the drug spread inside the thrombus over time and ensure the effectiveness of the drug and nanoparticles to remove the thrombus.

**Figure 2.**
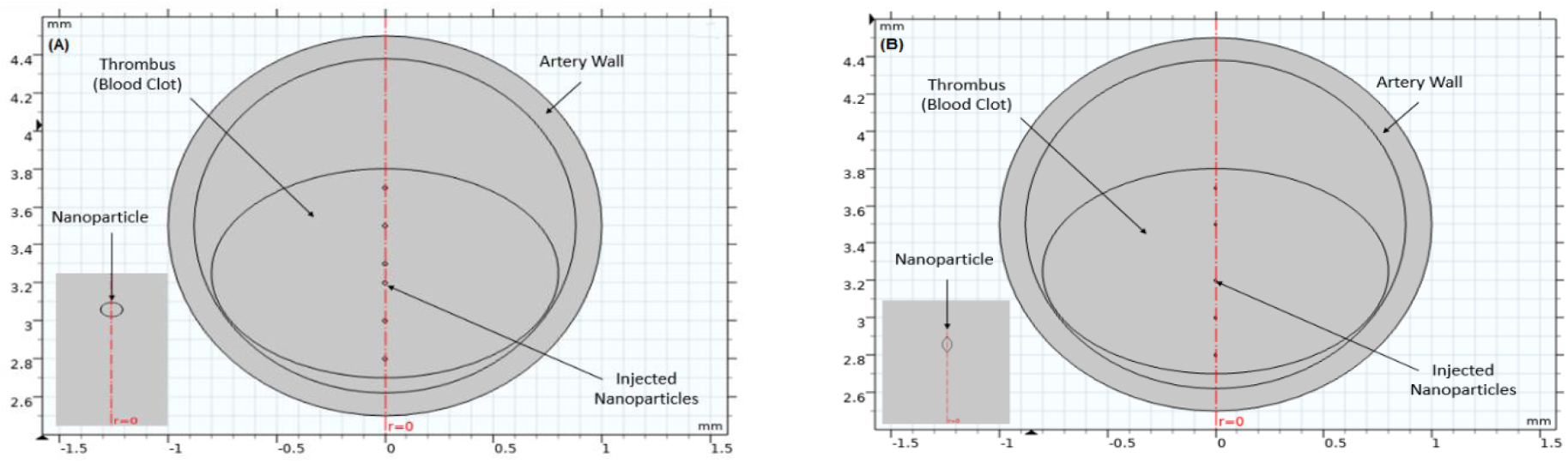
A scheme showing the 2-dimensional axial symmetrical geometry of the arterial vessel with a blood clot and injected polymeric nanoparticles with A) Sphere shape B) Multiply-twinned shape.

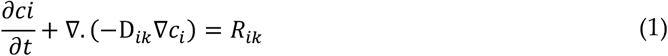

Where D_*ik*_ is the drugs i diffusion coefficient in the respective medium (m^2^/s) and *R_ik_* is the drugs i rate expression in the medium (mol/(m^3^·s)).

### 3.2 Moving Mesh and Convection-Diffusion Equation Modules

The diffusion equation was combined with the moving mesh node for modeling to simulate the thrombosis removal and shrinking during the time. The model consists of two-dimensional geometry that represents both thrombus, nanoparticles and artery as shown in (Figure 3). This clot field extends below the artery layer and under the shear pressure proportion that stimulates nanocarriers to start releasing the drug at the thrombus site and start dissolving and removing it.

**Figure 3.**
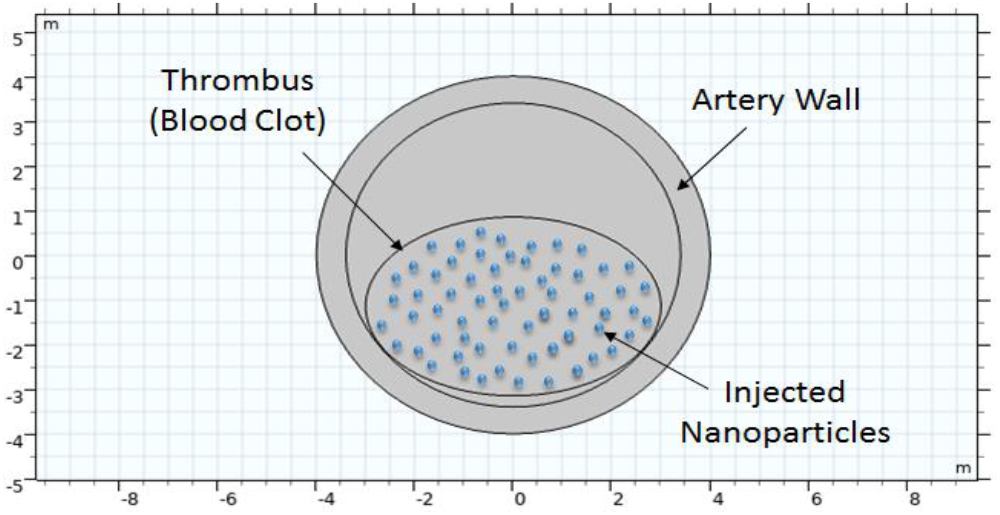
A scheme showing the 2-dimensional geometry of the arterial vessel with a blood clot and injected polymeric nanoparticles.

The materials of the model were selected automatically and the mesh was chosen to be a size (normal) mesh. The two models were also solved in the time domain to determine the thrombus response to the drug during the time after it was released as mentioned in the previous model.

## 3. Results and Discussion

When one of the veins or arteries becomes clogged as a result of blood clotting, then it must immediately intervene to treat this life-threatening problem. One of the most successful ways to do this task efficiently is to inject nanoparticles loaded with anti-clotting drugs, which immediately and within 3-5 hours dissolve the clot and prevent its growth without affecting the healthy vessels. The results of this study found significant and clear support for the efficacy and effectiveness of colloidal polymeric nanoparticles in dissolving and eliminating blood clots within a few hours. Also, in this section we discussed the modeling results of the drug diffusion and concentration inside the nanoparticle and the clots, besides the results of drug release from the polymer to the thrombus after exposure to shear pressure due to clotting, and the dissolve stages of this thrombus.

### 4.1 Chemical Reaction Engineering Module

In this model, after the polymeric nanoparticles have been injected to control the drug release location, this control allows to ensure the stability of the released amount of the drug from both of the sphere and multiply-twinned with prism-shapes nanoparticles during the time, it shouldn’t be in great quantities that lead to high toxicity, nor in small quantities that prevent the successful completion of the treatment and for the sake of patient’s safety.

Here the release of the drug has been studied for an hour and 30 minutes, and most often the time for dissolving the clot is from 2-4 hours. After the particles were injected into the coagulation site the drug was released at (0 min) time as shown in (Figure 4 A). But after these particles were subjected to shear pressure, the particles began to break down and a small amount of the drug was released in the 10th minute, as shown in (Figure 4 B).

**Figure 4.**
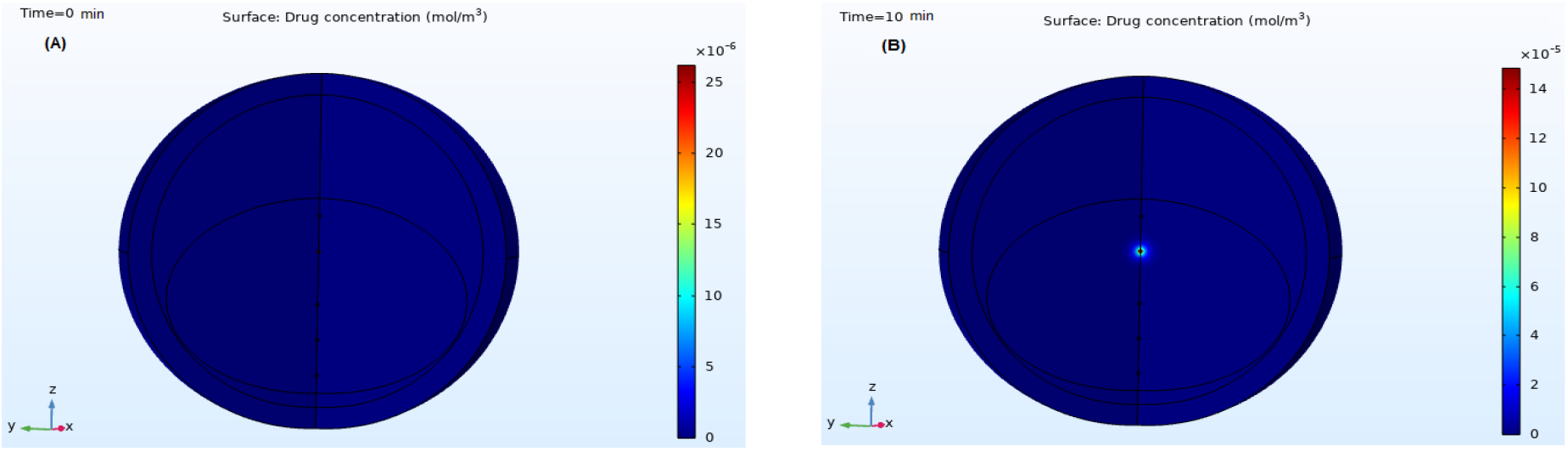
Drug diffusion and concentration of the sphere nanoparticle after A) 0 min and B) 10 min.

The Computational modeling adopted in this study is one of the new sources that provide a better understanding of the phenomena, behavior and the complex interactions between components of different designs for nanoparticles as carriers for medical drugs. It also provides a clear vision of the physical interaction between nanoparticles and thrombus behavior. Figure 5 provides visible and clear results of the anti-clotting drug release process from the multiply-twinned nanoparticle at the blood clot site at 0min and 10 min as shown in (Figure 5 A and B) respectively.

**Figure 5.**
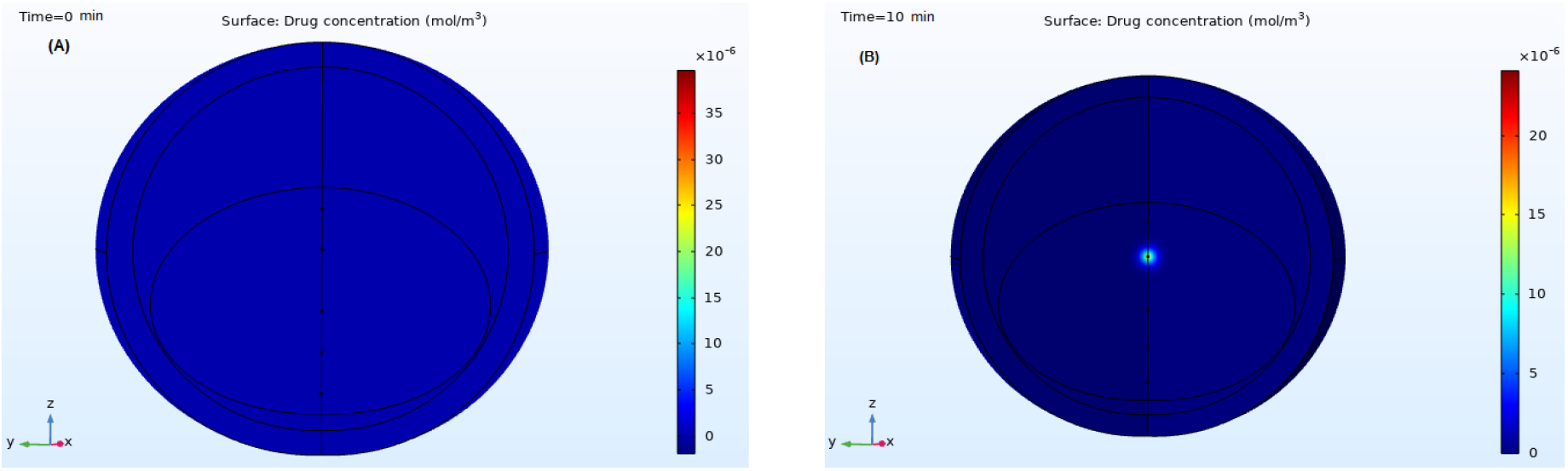
Drug diffusion and concentration of the multiple-twinned nanoparticle after A) 0 min and B) 10 min.

It is important to know that, both the amount of the drug-loaded into the nano-carriers, nanoparticles shape, decomposition of the encapsulated biomaterials, and the shear pressure intensity are the most important factors affecting the degradation of nanoparticles and the amount of released medicine. If we assume that the amount of both “Tissue Plasminogen Activator & Heparin” loaded in nanoparticle = 98% to be released fully in 4 hours and because the drug release is constant over time; the drug amount should be released in 1 hour from the total amount is 24.5%. As shown in (Figure 6 A) after 20 minutes, the amount of the drug release rate continues to increase steadily and after 50 minutes about 20% was released as shown in (Figure 6 B).

**Figure 6.**
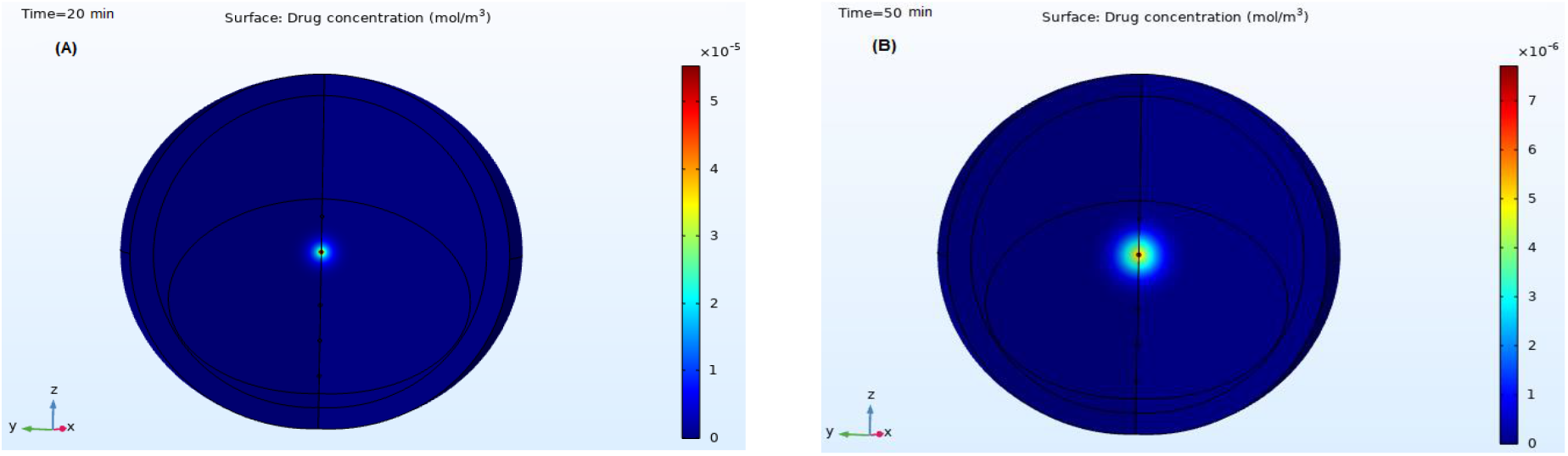
Drug diffusion and concentration of the sphere nanoparticle after A) 20 min B) 50 min.

Indeed, some of the therapeutic applications based on the nanoparticles require specific sizes and shapes to accurately control the drug released amount and time at the specified location in order to achieve the greatest therapeutic benefit. As cleared in (Figure 7 A and B) at 20th and 50th min respectively the diffused drug amount from the multiple-twinned nanoparticle is greater than the diffused drug amount from the sphere nanoparticle (Figure 6 A, B). To point out, the accurate control of the drug greatly contributes to anticipating the rates of active availability of the drug in the body, which is one of the basic conditions for the absorption of the therapeutic agent.

**Figure 7.**
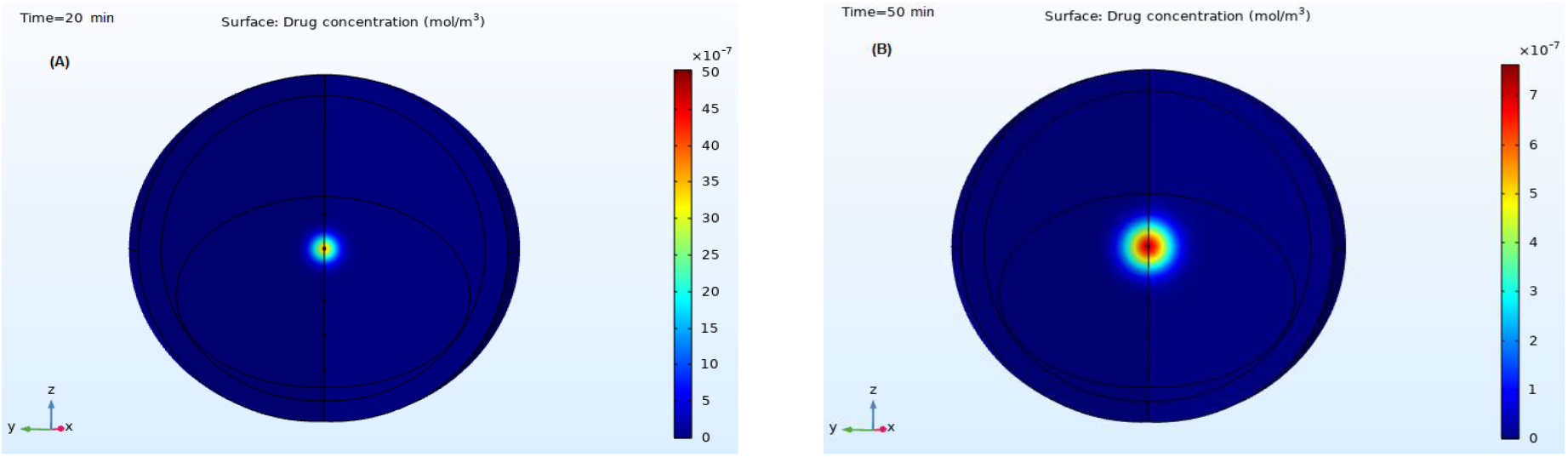
Drug diffusion and concentration of the multiple-twinned nanoparticle after A) 20 min B) 50 min.

In (Figure 8 A) in the 70th minute, 28% were fired, but in (Figure 8 B) the concentration of the drug is evident across the field of modeling as a result of the distributions of the concentration of all types involved in the drug release process as time functions, by dissolving the space-dependent mass balances in (Equation 1) on the 100th minute, 40% of the total drug was released.

**Figure 8.**
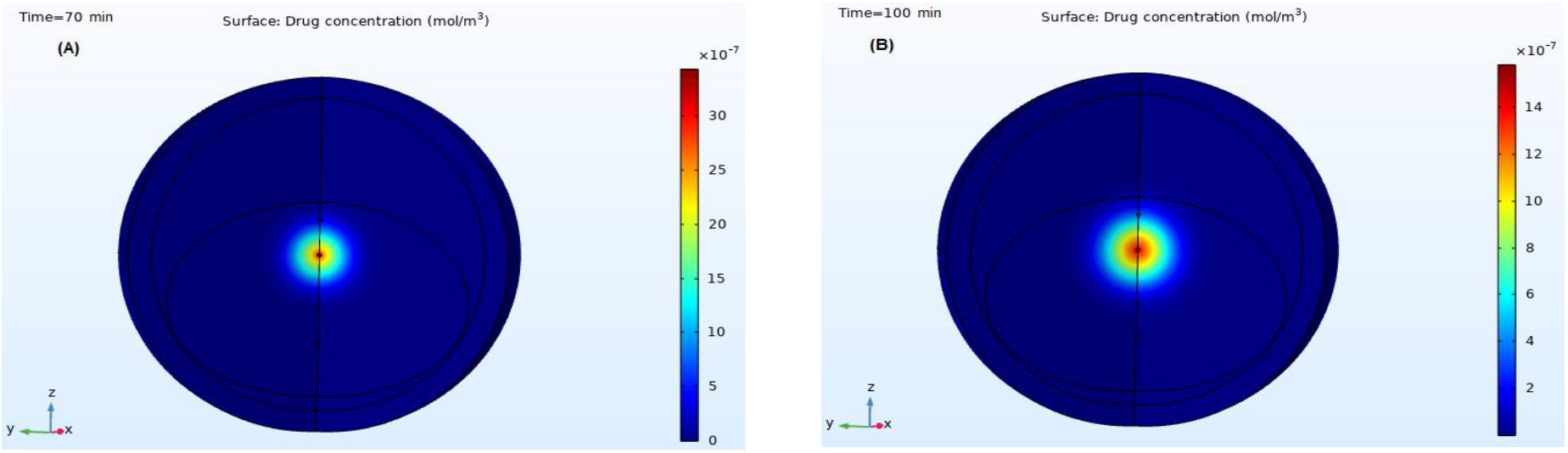
Drug diffusion and concentration of the sphere nanoparticle after A) 70 min B) 100 min.

To clarify, when developing drug delivery systems for therapeutic applications, it is very important to take into account both the drug biological decomposition in addition to the release profile for this drug which in turn contributes to improving and maximizing the therapeutic effect. The drug’s solubility and diffusion can accurately determine the effectiveness of these drugs as well as their active ingredients. As shown in (Figure 9 A, B), the nature of the release of drugs and anticoagulants from nanoparticles affected greatly by the nanostructure and geometry, and the drug that released in the 70th and 100th minute, respectively from the multiply-twinned shape showed that, the concentration of a drug that diffused outside of nanoparticles increases as a function of time with constant proportions.

**Figure 9.**
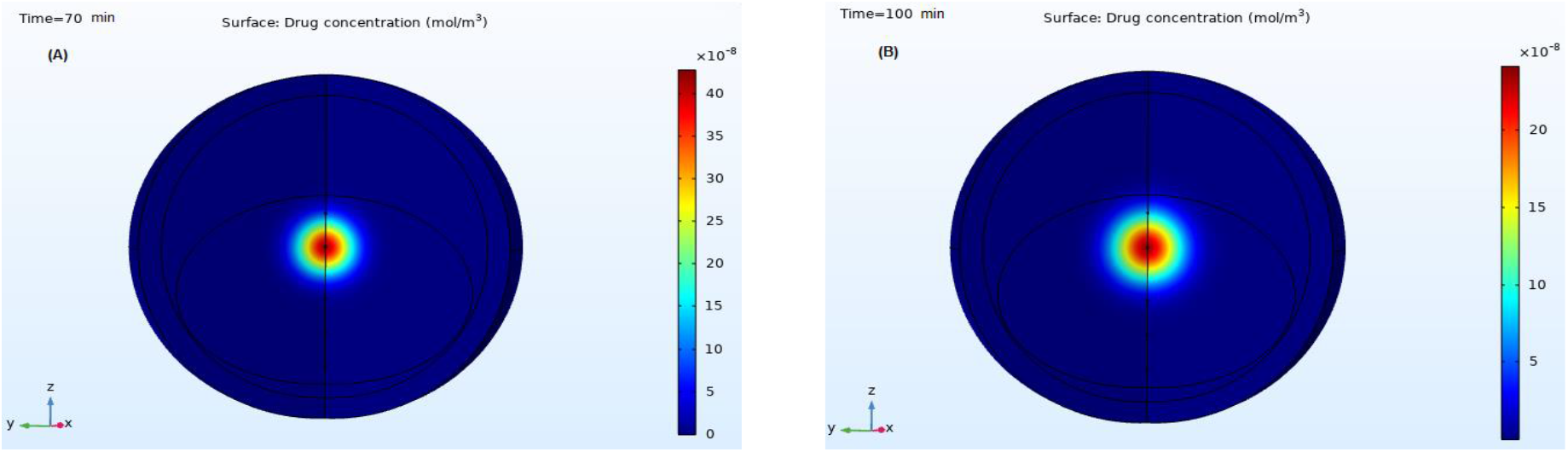
Drug diffusion and concentration of the multiple-twinned nanoparticle after A) 70 min B) 100 min.

As an illustration, the accurate design of nanoparticles greatly affects the place and time of the drug’s release due to the difference in sensitivity of these particles to body temperature, pH, pressure and enzymes. If we assume that, both of the two shapes adopted in the design of nanoparticles in this study release the same amount of medicine at the same time, the most important thing is to know the amount of the drug released is to know the concentration percentage for the diffused drug in the target site. The drug properties are largely controlled by both the shape and design of nanoparticles. The lower the amount of the medication that is released versus an increase in its concentration, the therapeutic efficacy of this drug will increase as well as the optimization of the patient’s compliance with this drug in less time. When both sphere and multiply-twinned nanoparticles shape are compared, the concentration of the drug in (Figure 10 B), which appears in red, is higher than the concentration of the drug in the spherical shape (Figure 10 A).

**Figure 10.**
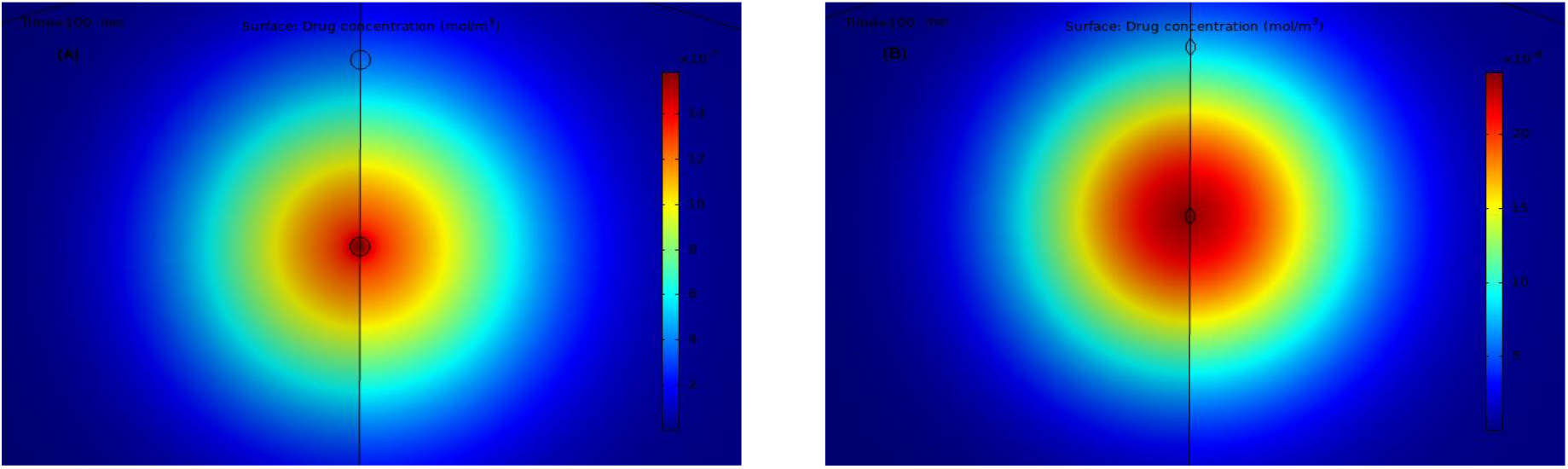
Drug diffusion and concentration after 100 min A) sphere shape B) multiple-twinned shape.

The concentration of the drug inside the nanoparticles depends on the particle shape and the time factor. Moreover, the drug concentration decreases with time inside the nanoparticles and increases overtime outside the nanoparticles due to the drug diffusion in the site of the thrombus and continues to be released in a regular manner and in constant quantities. Which facilitates the process of controlling the release of the drug at the site of the thrombus and obtaining the highest therapeutic efficacy. The color legend represents the concentration of the drug inside the nanoparticle so that the drug’s concentration increases (from blue to red). At (0 min) for the sphere shape the drug is concentrated inside the whole nanoparticle as shown in (Figure 11 A). As for the multiple-twinned nanoparticle the drug is concentrated in the core and gradually decreases as it gets closer to the shell as shown in (Figure 11 B).

**Figure 11.**
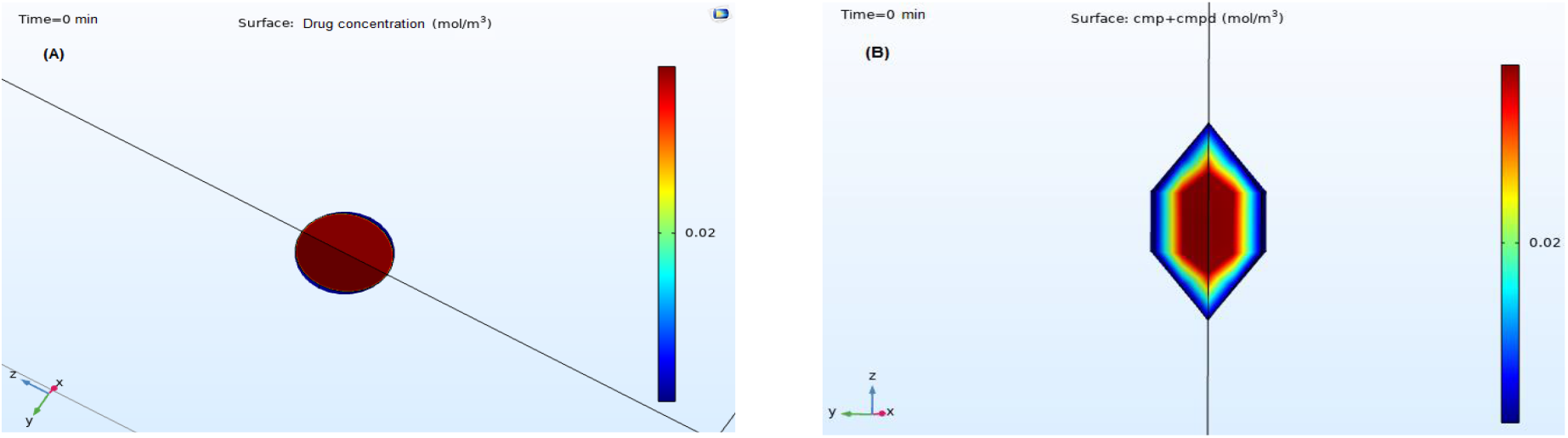
Drug concentration inside nano-carrier after 0 min A) sphere shape B) multiple-twinned shape.

After the first min (Figure 12 A) the magenta and blue colors become wider over time and the and red colors become narrow in the center over time because the drug has diffused outside the nanoparticles. In (Figure 12 B) the highest concentration of the drug is concentrated in the top and bottom of the twinned tip with red color.

**Figure 12.**
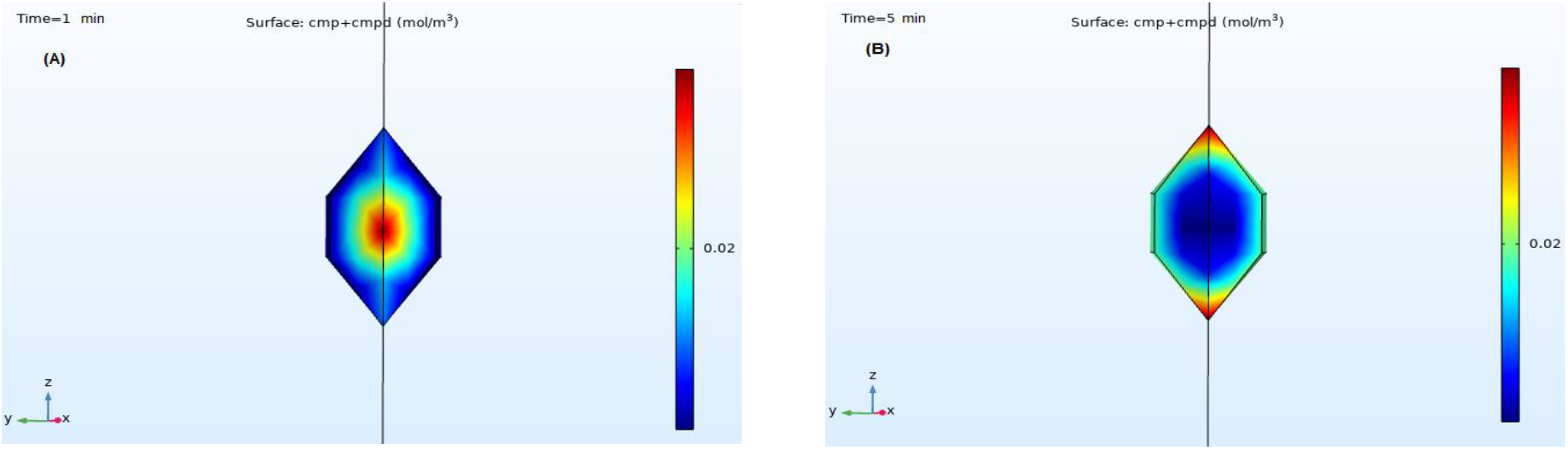
Drug concentration inside multiple-twinned nano-carrier after A) 1 min B) 5 min.

The magenta and blue colors index the low concentration of the drug and the red color shows the high concentration, and as shown in (Figure 13 A) in the 10th min the drug concentration inside nanoparticle was lowest in the center and highest in the inner surface because there are 94% of the drug still remains inside the nano-carrier. In the multiple-twinned shape, the drug is concentrated in the top and bottom of the twinned tip with red color as shown in (Figure 13 B) and remains constant inside the nanoparticle after the 10^th^ min.

**Figure 13.**
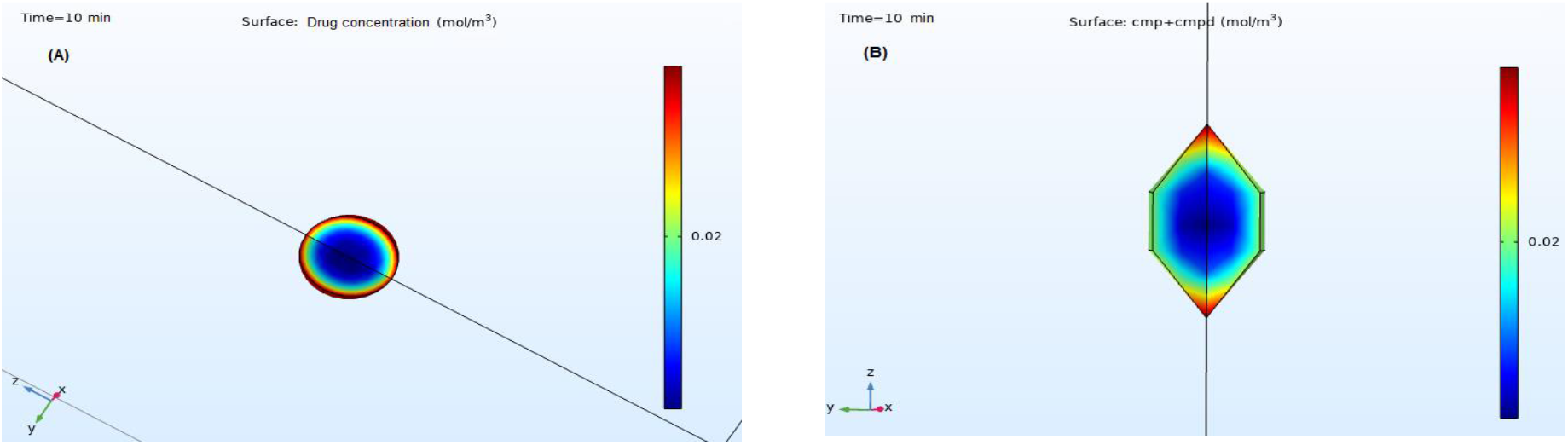
Drug concentration inside nano-carrier after 10 min A) sphere shape B) multiple-twinned shape.

After 100 min the release amount of the drug is about 40% and as shown in (Figure 14 A) the drug concentration inside the sphere nanoparticle becomes reduced after an hour and a half as the magenta and blue colors become narrow over time because the drug has diffused outside the nanoparticles. In an opposite manner in the multiple-twinned shape, the magenta and blue colors become wider over time and the highest concentration of the drug is concentrated in the top and bottom of the twinned tip with red color (Figure 14 B).

**Figure 14.**
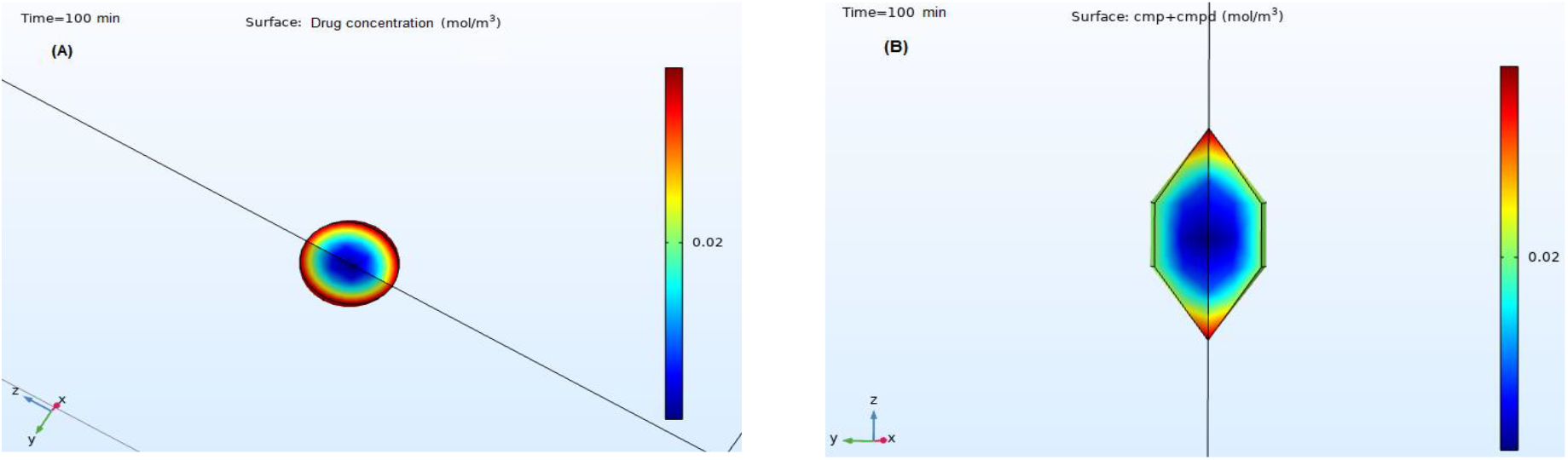
Drug concentration inside nano-carrier after 100 min A) sphere shape B) multiple-twinned shape.

Figure 15 describes the degradation of polymeric nanoparticles within an hour and a half as a result of the artery narrowing (Note: x-axis indicate the arc length (mm) and y-axis indicate the drug concentration of both Tissue Plasminogen Activator and Heparin (mol/m^3^)), stimulating the drug release at the thrombus site with a constant concentration over time as it begins to increase clearly during the 10th min from inside to outside the carriers (Figure 15 A) to reach the clot location and during the 100th min (Figure 15 B) blue carve showed that the concentration was the highest amongst other curves.

**Figure 15.**
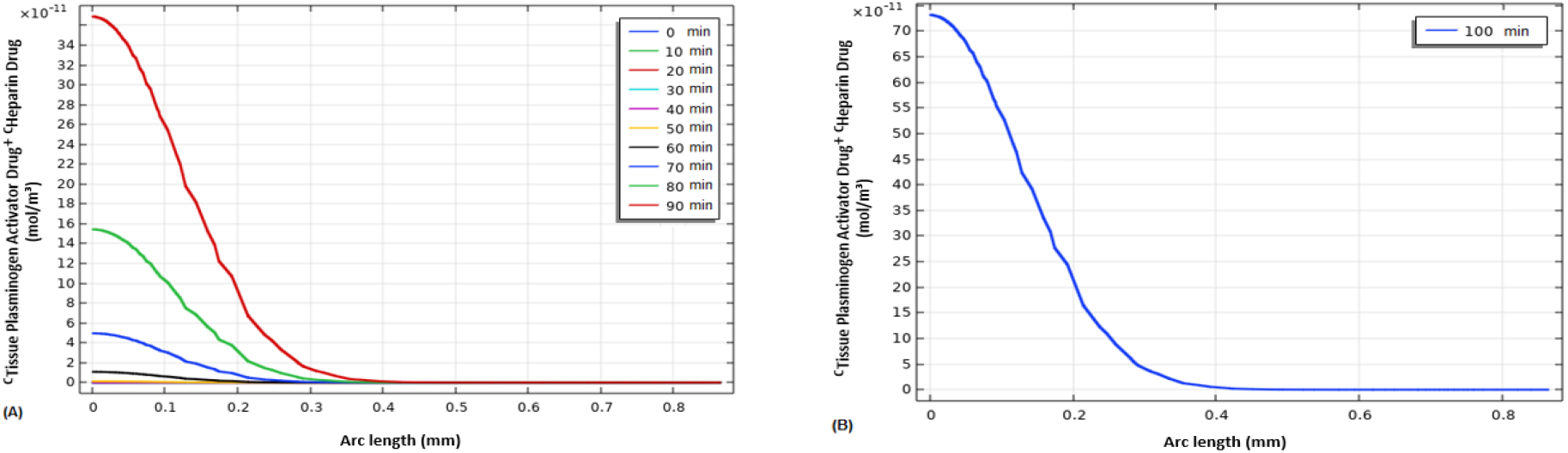
The total drug concentration profiles for the ‘Tissue Plasminogen Activator’ and ‘Heparin’ drugs in sphere nanoparticles at times up to A) 0-90 min B) 100 min across the modeling domain.

Figure 16 showed the total drug concentration profiles for the antithrombotic drugs in multiple-twinned nanoparticles and the drug concentration begin increased significantly from the 80^th^ min with the green curve and reach the highest in 100^th^ min with magenta carve and the Arc length remain constant in 0.4 mm.

**Figure 16.**
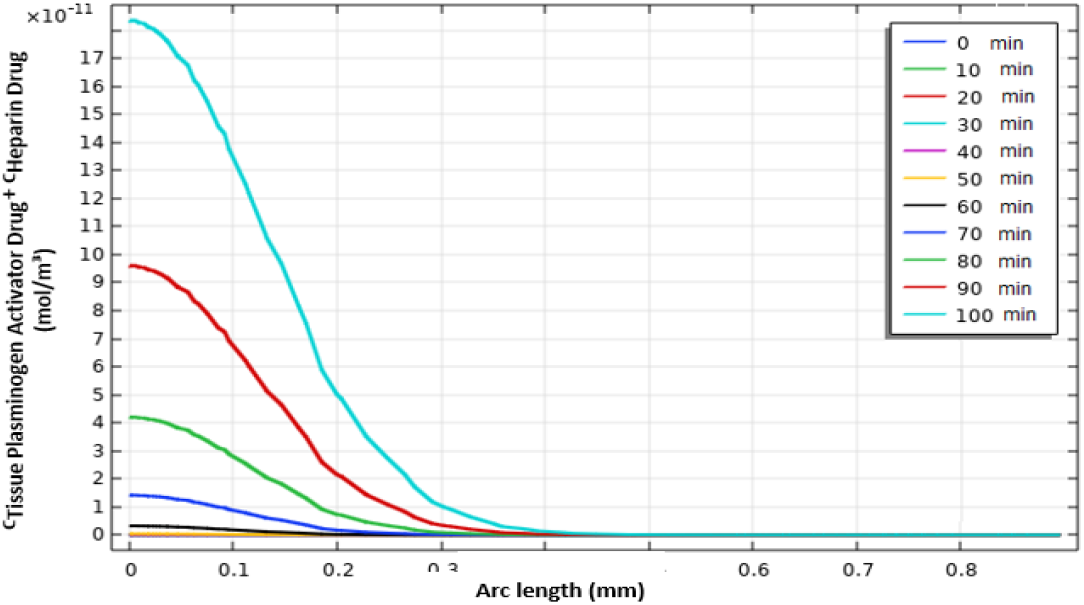
The total drug concentration profiles for the ‘Tissue Plasminogen Activator’ and ‘Heparin’ drugs in multiple-twinned nanoparticles at times up to 100 min across the modeling domain.

In fact, predicting the behavior of the drug release within the blood clot contributes to predicting the rate of change in the dimensions of the thrombus and the time of its dissolution. Markedly, the modeling of drug release from all nanoparticles gives a clearer understanding of the drug’s concentration inside the thrombus over time as shown in (Figure 17 A, B).

**Figure 17.**
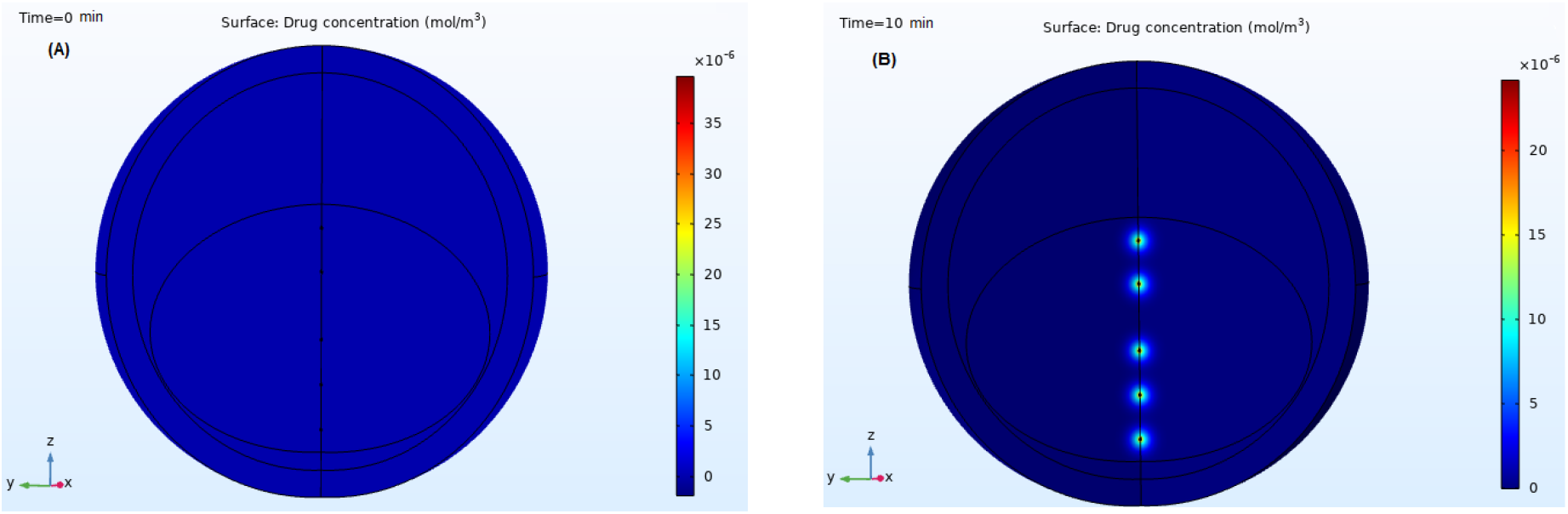
Drug diffusion and concentration from all of the multiple-twinned nanoparticles after A) 0 min B) 10 min.

The drug release profile from the injected nanoparticles becomes clear over time. If we imagine this model is 3D geometry definitely the Drug diffusion and concentration from all of the nanoparticles will be more visible inside the whole thrombus in view of the dispersion of nanoparticle. As shown in (Figure 18) the drug concentration in the 70^th^ and 100^th^ min is become the highest that appears in red color in the axial line which means the drug concentration after two hours will be the highest and in maximum which will reduce thrombus and reduce treatment time as will be explained in the next section of results.

**Figure 18.**
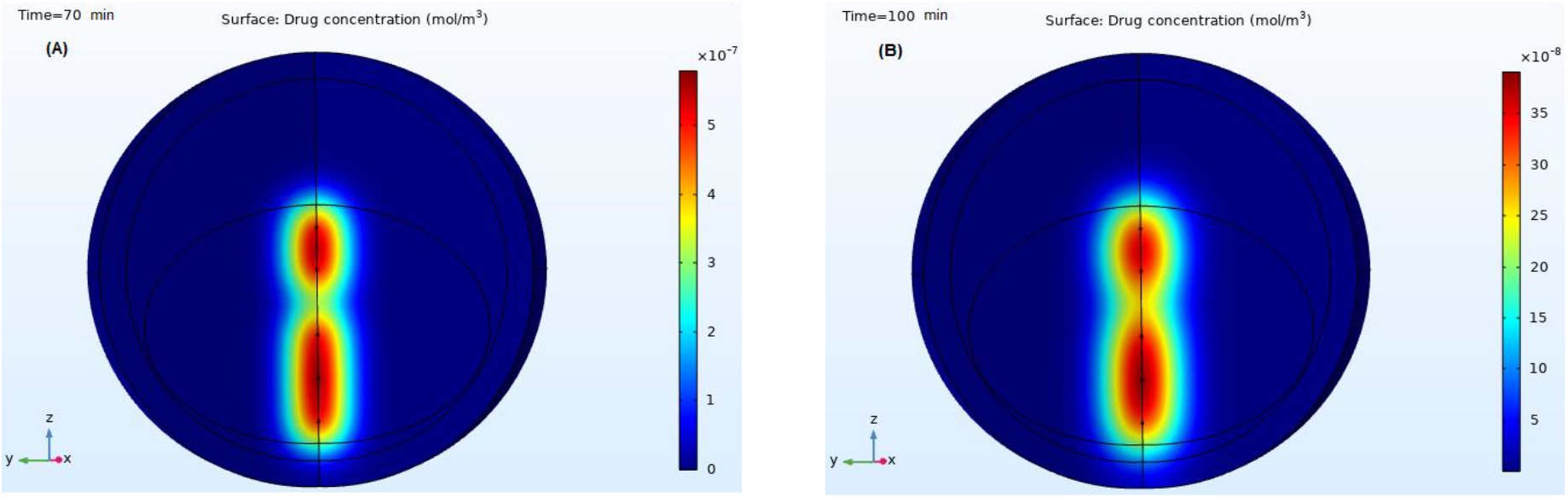
Drug diffusion and concentration from all of the multiple-twinned nanoparticles after A) 70 min B) 100 min.

### 4.2 Moving Mesh and Convection-Diffusion Equation Modules

The dissolving of the clots was simulated within an hour and a half from the release of the drug after injection of polymeric nanoparticles, which ranged in size from 0.1-0.3 nm. This model result is an important finding in the understanding of the blood clot elimination from the artery.

As shown in (Figure 19 A) in time (0 min) the blood clots have a smooth type with ≥ 6 mm thickness and area about 25-50% and 91°-180° circumference. in (Figure 19 B) in 20th min has a thickness of ≥ 5.5 mm and an area of about 50%.

**Figure 19.**
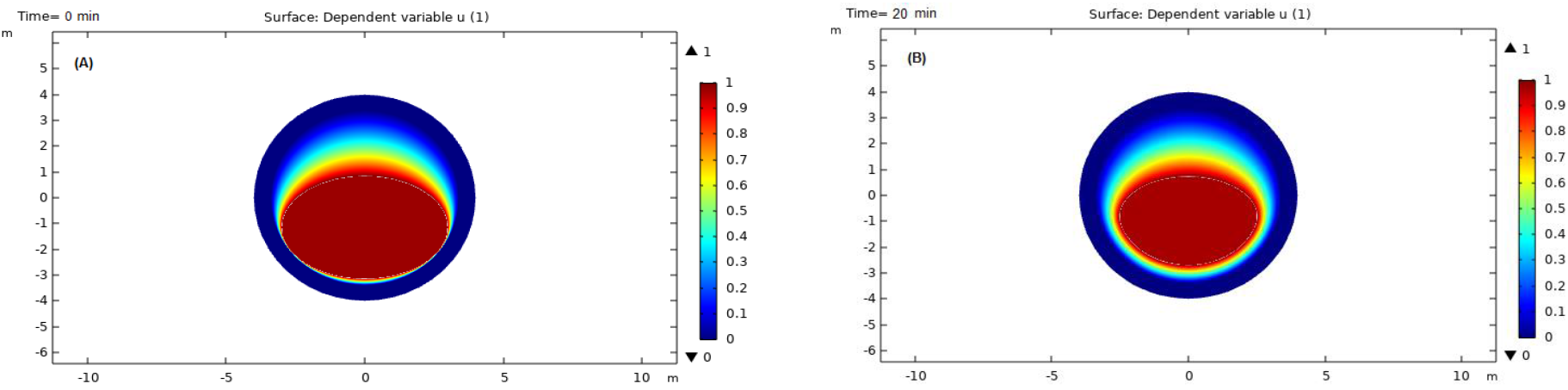
Blood clots dissolution after A) 0 min B) 20 min.

As shown in (Figure 20) the clots become smaller and small overtime in an acceptable manner which validates the nanoparticle’s efficiency and the anticoagulation drugs, in min 50th there were only clots remained with a thickness of ≥ 4.5 mm.

**Figure 20.**
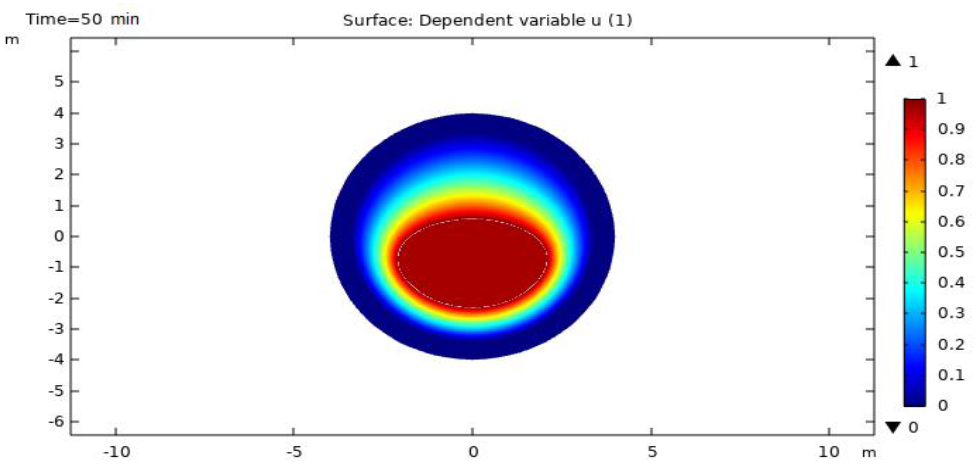
Blood clots dissolution after 50 min.

After 30 min later in the 80th min, there was only clots take less than 30% from the total channel area as shown in (Figure 21 A) and after 1:30 min the blood can flow in a good manner and this suggests that the 10% of the remained clots as shown in (Figure 21 B) can be totally removed after 2 hours. This result ties well with previous studies as it showed that, the polymeric nanoparticles can successfully treat for vascular diseases, especially thrombosis and strokes, that target the location of the blockage where these particles operate under the influence of shear pressure, ensuring that the drug will be released in the right place.

**Figure 21.**
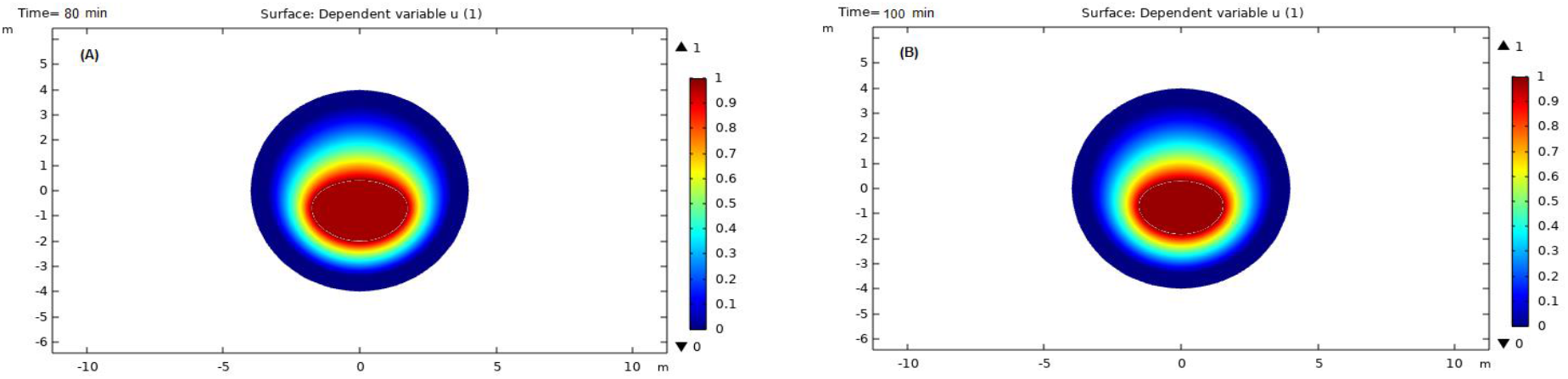
Blood clots dissolution after A) 80 min B) 100 min.

## 4. Conclusions

Nanotechnology has become increasingly used in medical applications because of its unique characteristics such as the surface to the volume ratio and its high capacity for drug loading, encapsulating efficiency and its ability in reducing side effects of traditional methods. Since blood clots are among the most dangerous life-threatening diseases, the basic findings of this study are consistent with emerging research showed that the effectiveness of nanotechnology in treating clots and saving many lives. In addition, these findings provide additional information about ‘Tissue Plasminogen Activator’ and ‘Heparin’ drugs in eliminating the coagulation for less than 4 hours. The chemical reaction engineering, moving mesh and convection-diffusion equation modules has shown its efficiency and accuracy in mimic the dissolution of the thrombosis, which these results can be admitted for the clinical trial applications and this may be considered a further validation of conducting further simulation using different nanoparticles types and another medication that are available for clot treatment. Finally, the nanoparticle shape plays an important role in drug delivery and efficiency due to the drug properties such as the loading capacity, distribution and in this study, the multiply-twinned shape was the best shape compared with the sphere as it showed the highest concentration of the drug in the thrombosis site.

